# CRISPR-Cas9 repair complexity in *Drosophila melanogaster*: NHEJ-induced deletions and HDR variability in the *bantam* microRNA gene

**DOI:** 10.1101/2025.05.29.656857

**Authors:** Pauline De Sousa, Elisabeth Houbron, Hervé Seitz, Isabelle Busseau

## Abstract

CRISPR-Cas9–mediated gene editing was used to generate specific mutants of the *bantam* gene in *Drosophila melanogaster*.

To drive non-homologous end joining (NHEJ) and achieve a precise deletion of most of the *bantam* locus, two guide RNAs targeting sites 90 base pairs apart were expressed in the germline using the UAS/GAL4 system. Thirty lethal and eight viable lines were established and analyzed. One lethal line exhibited the expected 90 bp deletion, while the others carried diverse indels at one or both cleavage sites. Among the viable lines, seven harbored a single-nucleotide deletion that did not disrupt *bantam* function. Notably, one viable line, *ban^d1-44^*, carried a hypomorphic allele that reduced organismal size without affecting viability.

To generate precisely edited *bantam* variants, CRISPR-Cas9–mediated homology-directed repair (HDR) was used using donor plasmids containing engineered mutations in the miRNA seed region, along with a scarless dsRED fluorescent marker. Approximately 40% of the resulting fluorescent lines were correctly edited, demonstrating the efficiency of this strategy for producing specific *bantam* variants. The remaining lines exhibited unexpected outcomes, among which partial HDR events, where the dsRED marker was integrated but not the *bantam* mutations, and full donor plasmid integrations, which led to duplication of the bantam locus.

These findings reveal the complexity of CRISPR-Cas9 outcomes, emphasizing the need for thorough screening and characterization of individual candidates in gene-editing experiments. They also provide valuable insights for optimizing genome editing strategies.

**Article summary:** The authors used Clustered Regularly Interspaced Short Palindromic Repeats (CRISPR) gene editing to mutate the *bantam* microRNA in *Drosophila melanogaster*.

They guided Non-Homologous End Joining to induce a precise 90 bp deletion. It was the rarest occurrence, while small inactivating indels occurred frequently.

They employed Homology-Directed Repair using a fluorescent marker to specifically target the *bantam* seed region. This efficiently produced the intended mutations but also led to unexpected outcomes, including partial sequence replacements and full donor plasmid integrations.

These results reveal the complexity of gene editing outcomes and highlight the importance of thorough molecular characterization in genome engineering experiments.

## INTRODUCTION

In 2012 the detailed characterization of the bacterial Clustered Regularly Interspaced Short Palindromic Repeats (CRISPR)-Cas9 system highlighted the possibility to adapt it for generating site-specific cuts in any genomic DNA (Jinek et al., 2012). The functioning of CRISPR-Cas9 system relies on the action of a double-strand nuclease, Cas9, guided by a specific RNA to target and cut DNA at a precise sequence. Once the DNA is cut, the cell activates its repair mechanisms that fix the break: either Non-Homologous End Joining (NHEJ) or Homology-Directed Repair (HDR). NHEJ directly ligates together the broken ends. It is a fast but imprecise mechanism, often leading to the production of insertions or deletions (indels). HDR is a precise repair mechanism that uses a homologous sequence as a template to repair DNA through recombination. It allows to generate specific genetic modifications in the target region, by introducing them into the template DNA.

The CRISPR-Cas9 system revolutionized genome editing as it can be adapted to any living organism besides being extensively deployed in model organisms. Pioneering studies in *Drosophila melanogaster* (Bassett et al., 2013; Gratz et al., 2013, 2014; Port et al., 2014; Ren et al., 2013) quickly established the best strategies and optimized usage parameters for both NHEJ and HDR (Bier et al. 2018; Zirin et al. 2022).

The *bantam* locus of *Drosophila* was first identified in a gain-of-function screen for genes that affect tissue growth (Hipfner et al., 2002). A 21 kb deletion representing the first loss of function allele *ban^Δ1^* was then generated. Animals homozygous for *ban^Δ1^* grew poorly and died massively during larval progression, occasionally producing small pupae which never hatched. All reported defects could be rescued using fragments of genomic DNA. These fragments were shown to express a microRNA hairpin precursor, demonstrating that the functional product of the *bantam* gene is a microRNA (Brennecke et al., 2003). Since, extensive studies have unveiled the essential function of *bantam* in tissue growth, both at the cell autonomous and at the systemic levels (Boulan et al., 2013). Many efforts were deployed over the past twenty years to identify *bantam* mRNA targets, including the use of computational tools such as TargetScanFly (Agarwal et al., 2018) which intends to predict miRNA targets on the basis of their perfect complementarity to the miRNA “seed” (nucleotides 2-7 of the mature miRNA), conserved among several insect species species. Experimental studies based on reporter constructs and/or biochemical associations highlighted candidate targets that respond to *bantam* at the molecular level. Although promising candidates have emerged from such approaches (Stark et al., 2003; Bassett et al., 2013; Banerjee and Roy, 2018; Hobin et al., 2022) those potentially directly implicated in growth control remain elusive.

To advance functional studies, the need is becoming compelling for mutations affecting *bantam* more precisely than the *ban^Δ1^* 21 Kb deletion. CRISPR-Cas9 is an ideal tool to achieve this. In a first part of the present work, we describe efforts to generate a precise deletion of the *bantam* hairpin precursor by mobilizing the NHEJ repair system. In the second part, we aimed at introducing precise modifications in the *bantam* seed in order to alter the specificity of target mRNA recognition, by providing a template engineered from the *bantam* locus. Both methods proved highly successful, generating much more edited animals than were needed. This high editing efficiency allowed a comprehensive characterization of many events, unveiling the variety of outcomes inherent to the process.

## MATERIALS AND METHODS

### *Drosophila* maintenance and stocks

Fly stocks were maintained under standard conditions at 25°C. Information on genes and sequence data were found in RRID:Flybase_FBst0473707 release FB2025_01 (Jenkins et al. 2022; Öztürk-Çolak et al. 2024).

The balancer stock *TM6B* carrying *Sb* and *Tb* homozygous lethal, recessive visible mutations (RRID:BDSC_23232) and the stocks carrying the *UASp-Cas9-P2* (RRID:BDSC_58985), *vasa-Cas9* (RRID:BDSC_55821), *nos-Cas9* (RRID:BDSC_54591) and *PiggyBac* transposase (RRID:BDSC_8285) transgenes were obtained from Bloomington Drosophila Stock Center (NIH funding P40 OD018537). The stock bearing the *nos-GAL4-VP16* driver was a gift from Pernille Rørth (Rørth 1998).

### *Drosophila* specific procedures

The single-fly DNA preps protocol for PCR was used on individual pupae or adults as described (Gloor et al., 1993).

Viability: the number of adults which developed from thirty L3 larvae in vials containing standard food was determined. Student t-test was performed, after verifying applicability conditions (Shapiro-Wilk wt p-value=0.1579; Shapiro-Wilk di44 p-value = 0.5679; Levene Pr(>F) = 0.745). Raw data are given in Table S1 in File S1.

Weight determination: five lots of 50 males per genotype were placed into pre-weighted 1.6 mL microtubes, then weighted on a Sartorius precision balance. Student t-test was performed after verifying applicability conditions (Shapiro-Wilk wt p-value=0.7443; Shapiro-Wilk d1-44 p-value = 0.4497; Levene Pr(>F) = 0.1562). Raw data are given in Table S2 in File S1.

Pupal size measurements: lots of eight pupae of each genotype were aligned on a plate, pictures were taken with a stereomicroscope, measures were done using ImageJ software, and volumes was given by the formula 4/3π(L/2)(w/2)2 (L, length; w, width) as in Boulan et al. 2013. Final pupal size values were shown as the ratio with respect to wild-type animals. An ANOVA test was performed on the log2-transformed data, after verifying applicability conditions (Shapiro-Wilk wt p-value=0.8397; Shapiro-Wilk d1-44 p-value=0.1765; Shapiro-Wilk dΔ1 p-value=0.9688; Shapiro-Wilk d1-44/dΔ1 p-value=0.2933; Levene Pr(>F) = 0.07052). After demonstrating a significant difference between at least two groups, a post-hoc Tukey test was conducted. Raw data are given in Table S3 in File S1.

### General molecular biology procedures and plasmid construction

Depending on the expected fragment size, the GoTaq® G2 DNA Polymerase (Promega) or the Q5 High-Fidelity DNA Polymerase (New England Biolabs) were used according to the supplier’s recommendations. Preparative PCR for plasmid constructions were systematically performed with Q5 High-Fidelity DNA Polymerase. The resulting fragments were checked on 1 or 2 % agarose gels and purified using the QIAquick PCR Purification Kit (Qiagen) when needed. The sequences of oligonucleotides used in the study are given in Table S4 in File S1. DNA sequencing services were provided by eurofinsgenomics.eu (Sanger) and plasmisaurus.com or smartlifebiosciences.com (long read). Qiagen Plasmid Kits were used to prepare high quality plasmid DNA. All plasmid constructs were verified by long run sequencing. *Their annotated sequences will be available at* https://github.com *– in progress*.

Guide RNAs with no predicted off-targets in the *Drosophila melanogaster* genome available at the time (dm6 assembly) were identified and evaluated using CRISPR Target Finder (RRID:SCR_023641 http://targetfinder.flycrispr.neuro.brown.edu/). Their sequences were integrated into vectors pCFD5 and pCFD6 (gifts from Simon Bullock, RRID:Addgene_73914 and RRID:Addgene_73915, respectively), following the protocol described in www.crisprflydesign.org (Port and Bullock 2016) except that the NEBuilder® HiFi DNA Assembly Master Mix was used instead of Gibson Assembly.

*pCFD6-bansg58* for NHEJ was designed to express two small guide RNAs targeting the *bantam* locus. A transgenic strain was produced by The BestGene service (https://www.thebestgene.com RRID:SCR_012605) using the PhiC31 system into strain *y1 w67c23; P{CaryP}attP2 (67A)* (RRID:BDSC_8622). A stock carrying both *UASp-Cas9-P2* and *PCFD6-bansg58* transgenes was established following genetic crosses and recombination between the two transgenic lines.

*pCFD5-sgeban7* for HDR was designed to express two small guide RNAs, one targeting the *ebony* gene (initially to facilitate screening) and the other one targeting the *bantam* locus.

Donor plasmids for HDR using the scarless method (http://flycrispr.molbio.wisc.edu/scarless and Tindell et al., 2020) derive from a pBT-*bantam* construct containing 3256 bp encompassing the *bantam* locus inserted in pBluescript as described earlier (Busseau et al. 2024). Other recombinant plasmids were generated using the NEBuilder® HiFi DNA Assembly Master Mix following the supplier’s instructions. To produce pBT-*bantam*-dsRED, a 6134 bp PCR fragment was amplified from pBT-*bantam* using primers d2729 and d2731, and assembled with a 1715 bp fragment amplified using primers d2728 and d2730 from pScarlessHD-DsRed, a gift from Kate O’Connor-Giles (RRID:Addgene_64703). This resulted in the disruption of the Protospacer adjacent motif (PAM) by the *PiggyBac-3xP3*-dsRED cassette insertion. Then, a 7779 bp fragment was amplified from pBT-*bantam*-dsRED using primers d2436 and d2437, and assembled with annealed oligonucleotides designed to replace the wild-type *bantam* sequences with mutated ones : d2674 and d2675 to produce pBT-*ban^m11^-dsRED*, d2678 and d2679 to produce pBT-*ban^m13^-dsRED*, d2682 and d2683 to produce pBT-*ban^m15^-dsRED*.

### CRISPR-Cas9-NHEJ experiments and analysis

Virgin females bearing the germline driver *nos-GAL4-VP16* were mated to double transgenic *UASp-Cas9-P2* ; *PCFD6-bansg58* males at 25 °C or 29 °C (Table S5 in File S1) to produce offsprings expressing Cas9 and the two guide RNAs in the germline. These were then crossed to the *TM6B* balancer stock carrying *Sb* and *Tb* mutations, and individual males from their progeny were used to establish lethal or viable stable lines.

The molecular organization of the *bantam* locus in edited lines was determined by Sanger sequencing of PCR fragments amplified using d2014 or d1882 and d2016 (expected size of 542 bp or 499 bp, respectively, in the wild type).

### CRISPR-Cas9-HDR experiments and analysis

For each *bantam* variant to be edited, mixtures of 300 ng/µL of *pCFD5-sgeban7* with 500 ng/µL of donor plasmid were injected into 400 *vasa-Cas9* embryos by the *Drosophila* transgenesis facility of the CBM (https://www.cbm.uam.es/). Adult survivors were recovered and individually crossed to 4-5 flies from the TM6B balancer stock and their progeny was screened for fluorescent eyes. Out of 400 embryos injected with the donor plasmids pBT-*ban^m11^-dsRED*, pBT-*ban^m13^-dsRED*, and pBT-*ban^m15^-dsRED,* 100, 191, and 87 fertile adults survived, respectively. Among these, 5, 13, and 12 individuals produced at least one fluorescent progeny (Table 1).

**Table 1:**
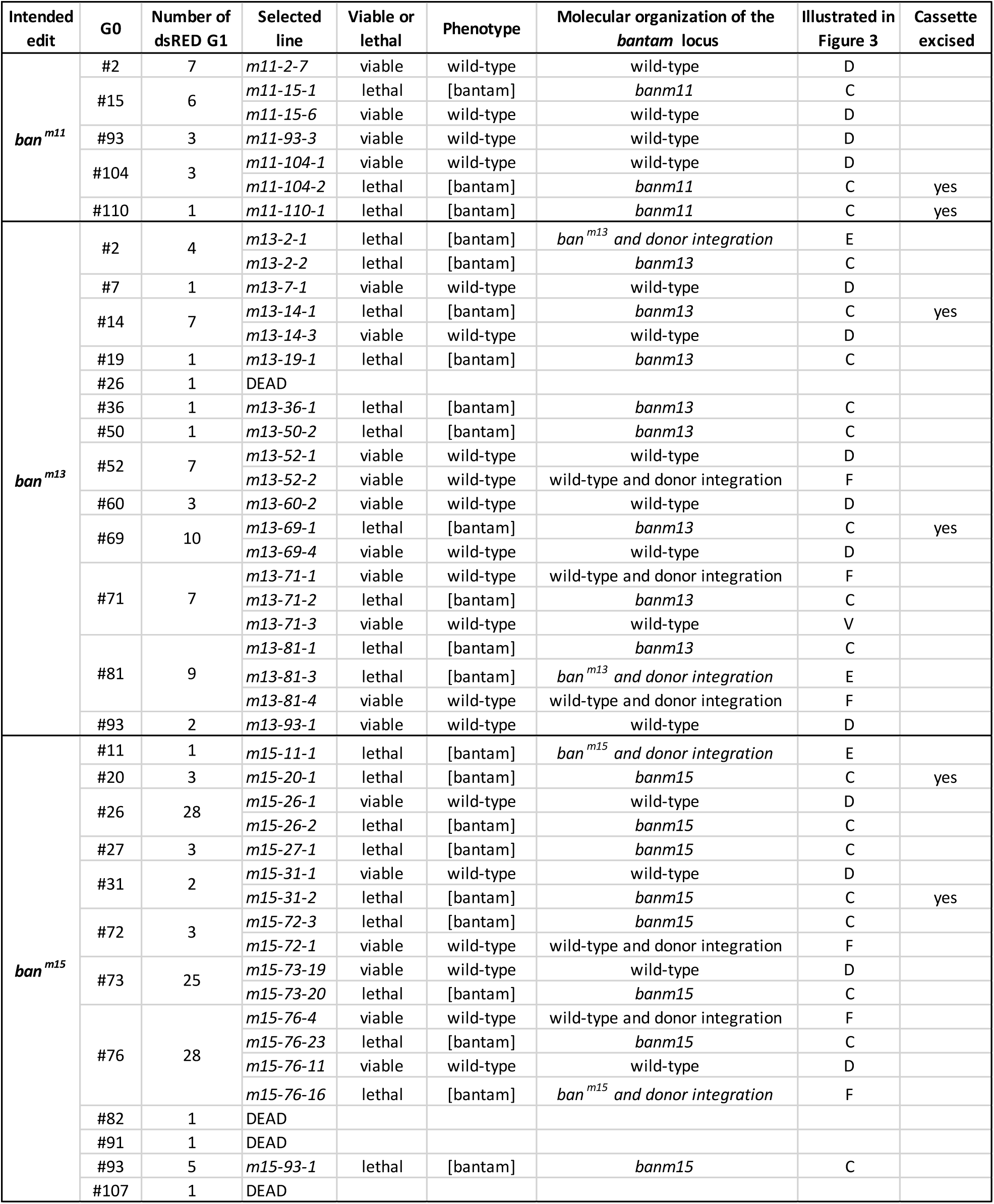
Lines generated by CRISPR-Cas9-HDR.

Long-range PCR to amplify the *bantam* region were performed using oligonucleotides d1299 and d1314 (expected size 4 kb in the wild-type). PCR to amplify the junction between pBluescript and the right homology arm were performed using oligonucleotides d1605 and d1313 (expected size 489 bp). The two parts of the duplication of the *bantam* region were amplified separately using oligonucleotides d1299 and d1605 (proximal part) and d1609 and d1314 (distal part).

In addition, in the case of *ban^m15-73-17^* a large fragment of approximately 14 Kpb containing the entire duplicated region including the pBluescript vector was amplified between oligonucleotides d2905 and d2906 designed to support NEBuilder® HiFi DNA Assembly. The resulting plasmid was transformed into bacteria, extracted and subjected to long read sequencing (Figure S1).

Precise excision of the *PiggyBac-3xP3*-dsRED cassette from selected lines was done by crossing into PiggyBac Transposase background and screening for fluorescence loss as described (Tindell et al., 2020).

## RESULTS

### CRISPR-Cas9-NHEJ to create deletions in the *bantam* locus

To create deletion mutants of the *bantam* gene, Cas9 and two sgRNAs targeting two sites 90 bp apart on each side of *bantam* were expressed in the germline. This was expected to lead to the removal of the functional part of the *bantam* locus after NHEJ repair, and thus to create a complete loss of function mutation (Figure 1A). As complete loss-of-function mutations of *bantam* have been reported to cause lethality at the pupal stage (Brennecke et al., 2003, Boulan et al., 2013), stable lines were established from the progeny using a TM6B third chromosome balancer stock marked with *Sb* and *Tb*. Of the thirty-six lines tentatively established, twenty-eight maintained the balancer chromosome over successive generations (more than forty so far), indicating that they were homozygous lethal, while eight could be maintained as homozygous stocks in the absence of the balancer chromosome, indicating that they were homozygous viable.

**Figure 1:**
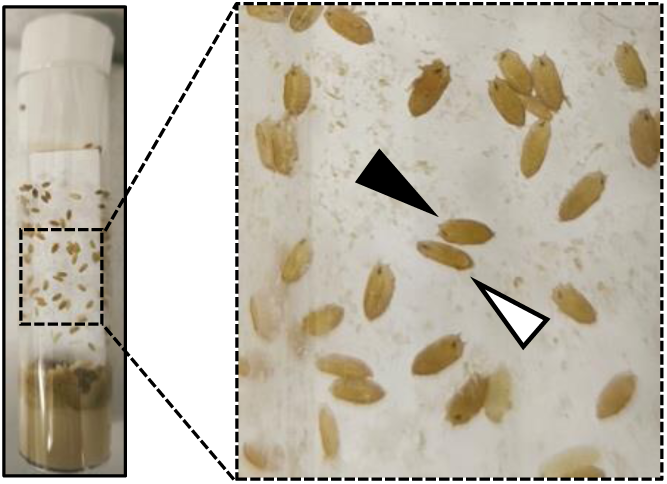
Typical appearance of a vial containing a *bantam* loss-of-function mutant stock. Open arrowhead points to a pupa homozygous for a *bantam* loss of function mutation (here as an example *ban^d1-67^*) mutation. Plain arrowhead points to a pupa heterozygous over *TM6B Tb Sb*.

The molecular structure of the *bantam* locus was subsequently determined in homozygous animals. For the eight viable lines, homozygous adults were examined. For the twenty-eight lethal lines, we took advantage of previous findings indicating that *bantam* homozygotes die at the pupal stage to identify homozygous pupae based on the absence of the *Tb* marker. Using this approach, we were able to analyse twenty-five of the twenty-eight homozygous lethal lines. The remaining three lines were excluded due to the failure to recover any Tb-negative pupae across three generations.

We noticed that homozygous pupae produced by the twenty-five lethal lines that we studied looked markedly smaller than wild-type pupae (Figure 1), consistent with published *bantam* loss of function phenotype (Brennecke et al., 2003; Boulan et al., 2013).

Homozygous animals of edited lines were individually processed for Sanger sequencing of the *bantam* region. The results are shown in Figure 2 and File S2.

**Figure 2:**
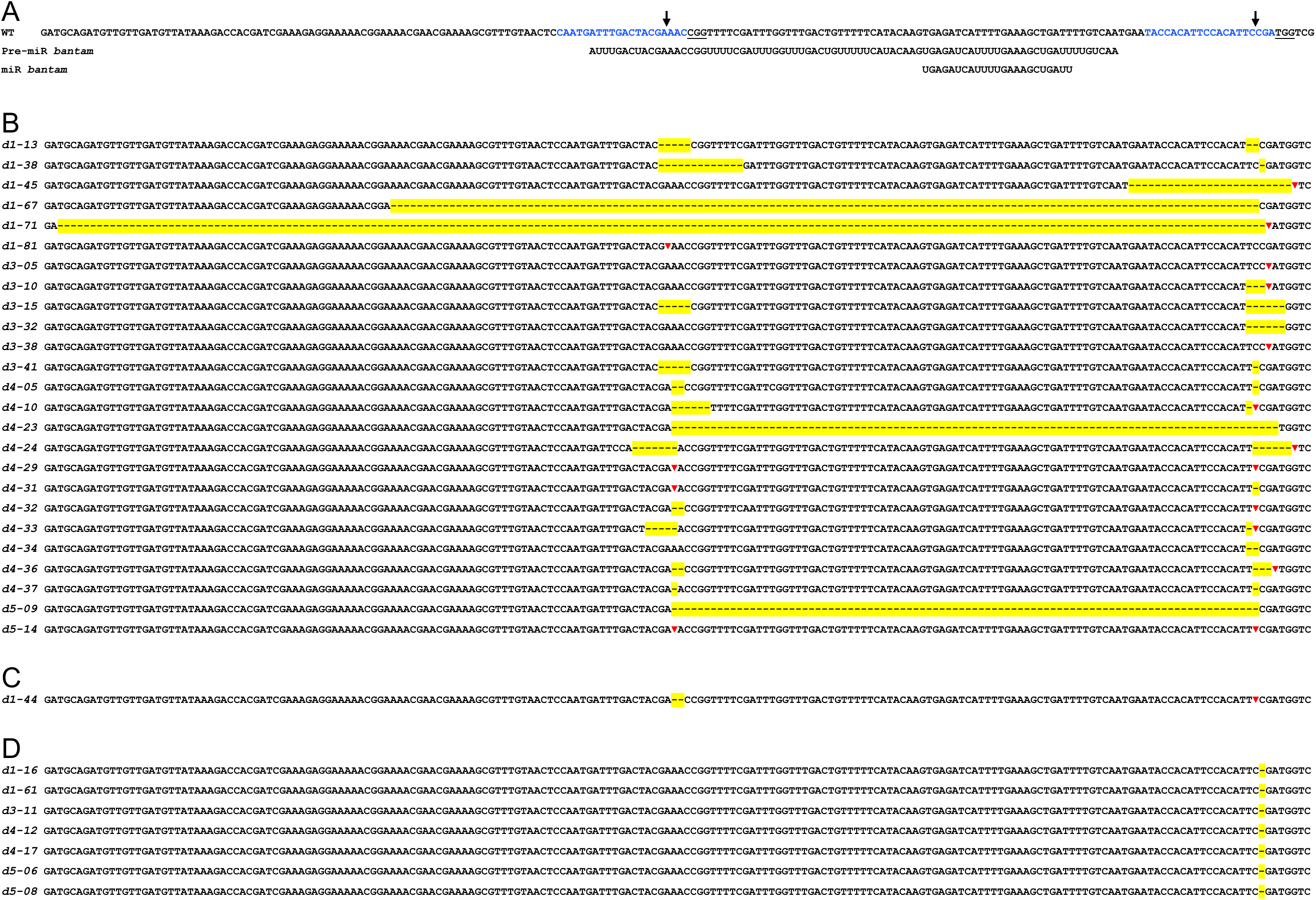
Alleles of *bantam* generated by CRISPR/Cas9/NHEJ. **A.** Genomic sequence of the wild type *bantam* locus showing the target sequences of two guide RNAs (blue) and the protospacer adjacent motives (underlined). The predicted positions of cleavage sites are indicated by a vertical arrow. The pre-miR and the mature *bantam* RNA sequences are shown below the genomic sequence. **B.** Genomic organization of the edited *bantam* locus in homozygous lethal lines. Deletions are represented with dashes highlighted in yellow, positions of insertions are indicated by red triangles. Full sequences are in Figure S1. **C.** Genomic organization of the *bantam* locus containing the *ban^d1-44^* mutation. **D.** Genomic organization of the edited *bantam* locus in homozygous viable lines.

The twenty-five lethal lines that were sequenced displayed some indels around one or the other or both predicted Cas9 cleavage sites. Only one of them, *ban^d5-09^*, had a large deletion of 90 bp exactly between the two cleavage sites. Three had large deletions extending beyond one or both cleavage sites: *ban^d4-23^* (deletion of 93 bp), *ban^d1-67^* (deletion of 135 bp), and *ban^d1-71^* (deletion of 186 bp). The other 21 lines had indels at the vicinity of either the distal (one case, *ban^d1-81^*, 11 bp insertion) or the proximal site (six cases), or both (fourteen cases). Around half of the mutants showed clear cut deletions, sometimes as small as a single nucleotide (*ban^d4-37^*). The rest had insertions of very variable sizes, from two (*ban^d3-38^*, *ban^d4-24^*) to ninety (*ban^d4-29^*) nucleotides, often concomitant to a deletion. In many cases the inserted sequences could easily be traced from the *bantam* locus itself, generating a duplication (exampled underlined in Figure S1).

None of the eight viable lines contained the intact wild-type sequence. Seven of them, despite being phenotypically wild type, displayed a single cytosine nucleotide excision at the distal Cas9 predicted cleavage site. This polymorphism was absent from the parental strains used to generate them (transgenic *nos-GAL4-VP16*, *UAS-Cas9-P2, and pCFD6-bansg58*) (Figure S1), and therefore most likely arose during NHEJ repair.

Finally, one event only partly affected *bantam* function, producing the viable, hypomorphic allele *ban^d1-44^* that is described below and in Figure 3.

**Figure 3:**
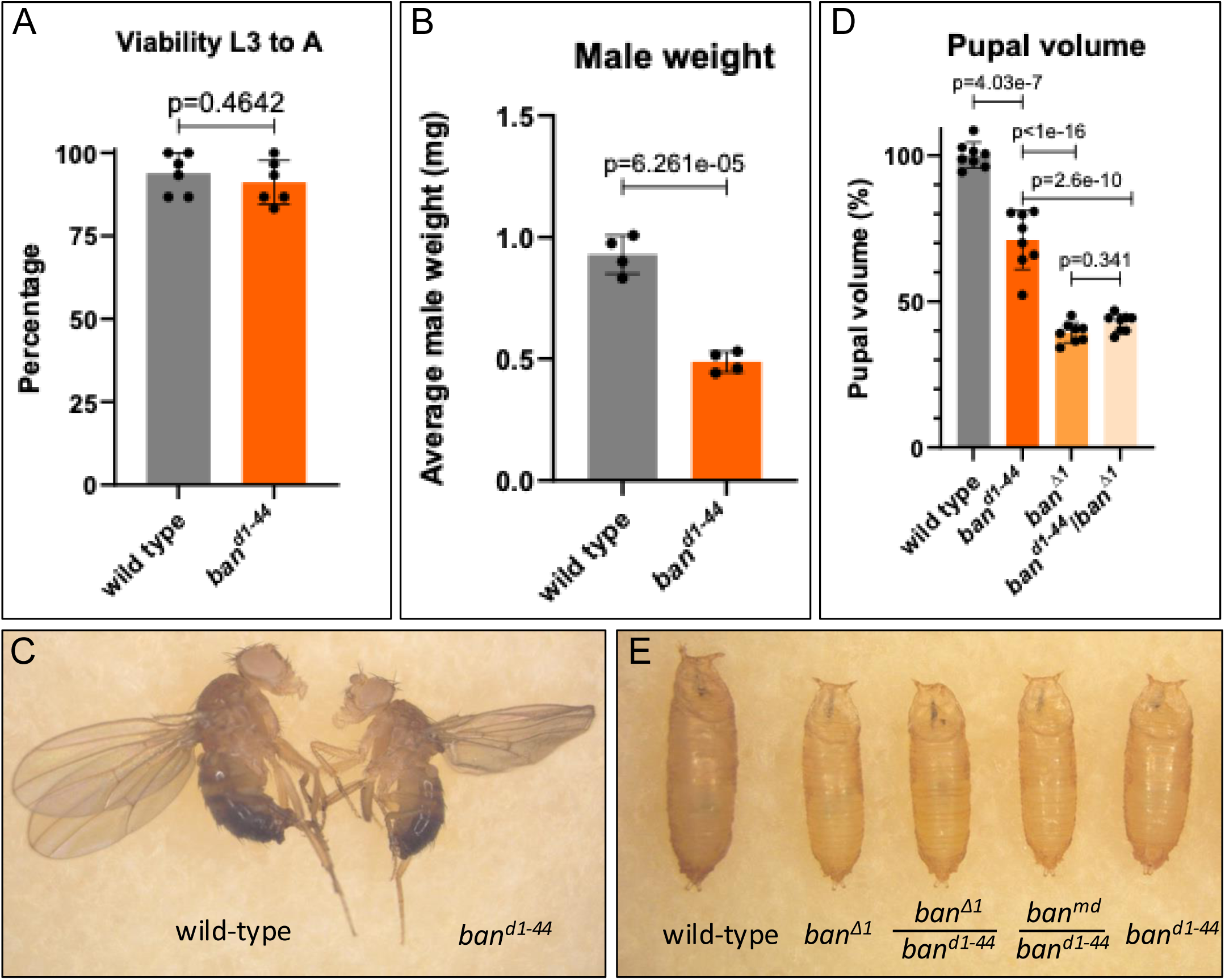
Hypomorphic allele *ban^d1-44^*. **A.** Viability of wild-type and *ban^d1-44^* homozygous third-instar larvae (six batches of thirty animals) to the adult stage. Relevant p-values (p) are indicated. **B.** Average weight of a wild-type male and a *ban^d1-44^* homozygous male, inferred from the total weight of four batches of fifty males per genotype. Relevant p-values (p) are indicated. **C.** Representative examples of wild-type and *ban^d1-44^* homozygous adult males. **D** Volumes of *bantam* mutant pupae expressed as percentages of the wild-type average (n=8 for each genotype). Relevant p-values (p) are indicated. **E.** Representative examples of wild-type and *bantam* mutant pupae.

In conclusion, the CRISPR-Cas9-NHEJ system using two sgRNA 90 bp apart, initially intended to generate precise deletion alleles of *bantam,* did so in a single case (*ban^d5-90^*) out of more than thirty events that were characterized at the molecular level. Still, the majority of CRISPR-induced rearrangements did inactivate *bantam*, either by creating large deletions encompassing the two predicted Cas9 target sites or by generating small indels at one or both cleavage sites. A non negligeable number of events altered the *bantam* sequence in a phenotypically inconsequential way.

### A novel hypomorphic allele of *bantam*

The *ban^d1-44^* line captured our attention because, despite being fully viable (Figure 3A), homozygous *ban^d1-44^* flies weighed only half as much as wild-type flies and exhibited a visibly reduced size (Figure 3B, C). Besides, the volumes of homozygous *ban^d1-44^* pupae were reduced by 30% relative to wild-type pupae (Figure 3D, E). The adults that emerged from these small pupae were small but healthy and fertile. Notably, homozygous *ban^d1-44^*pupae were not as drastically reduced in size as homozygous *ban^Δ1^* null mutant pupae, which appeared thinner and exhibited a 60% reduction in volume compared to wild-type pupae.

Complementation tests with the original *ban^Δ1^* deletion allele (Hipfner et al., 2002) and a recently characterized *ban^md^*null mutation (Busseau et al., 2024) confirmed that the *ban^d1-44^*mutation is a hypomorphic allele of *bantam* (3E). Hemizygous *ban^d1-44^/ban^Δ1^*or trans-heterozygous *ban^d1-44^/ban^md^* pupae displayed phenotypes indistinguished from that of homozygous *bantam* null *ban^Δ1^* mutants, including lethality.

At the molecular level, the *ban^d1-44^* line did not appear qualitatively different from many other lines that were homozygous lethal. It carries a small deletion of two nucleotides at the proximal cleavage site and an insertion of twenty-five nucleotides at the distal cleavage site. This insertion comprises two motives CACATTCCACA and ATACACATTCCA, which seem to be duplicated from the sequence ATACCACATTCCACA adjacent to the cleavage site (Figure S1).

The *ban^d1-44^*hypomorphic allele provides a unique tool to uncouple the growth defects associated with *bantam* loss-of-function mutations from their lethality, making it highly valuable for future functional studies of *bantam*.

### CRISPR-Cas9-HDR to generate precise *bantam* variants

To generate *bantam* gene variants with altered target specificities, mutations affecting the seed region of the miRNA (Figure 4A) were introduced into donor plasmids designed for scarless allele replacement. These donor constructs included a *PiggyBac-3xP3*-dsRED cassette to facilitate the identification of fluorescent-eyed animals (Figure 4B). The mutations *ban^m11^*, *ban^m13^*, and *ban^m15^* were engineered to alter the miRNA seed sequence while preserving the predicted secondary structure of the precursor RNA (Busseau et al., 2024). Donor plasmids were co-injected with a guide RNA-expressing vector into embryos expressing germline Cas9, and fluorescent (dsRED-positive) progeny were used to establish stable lines.

**Figure 4:**
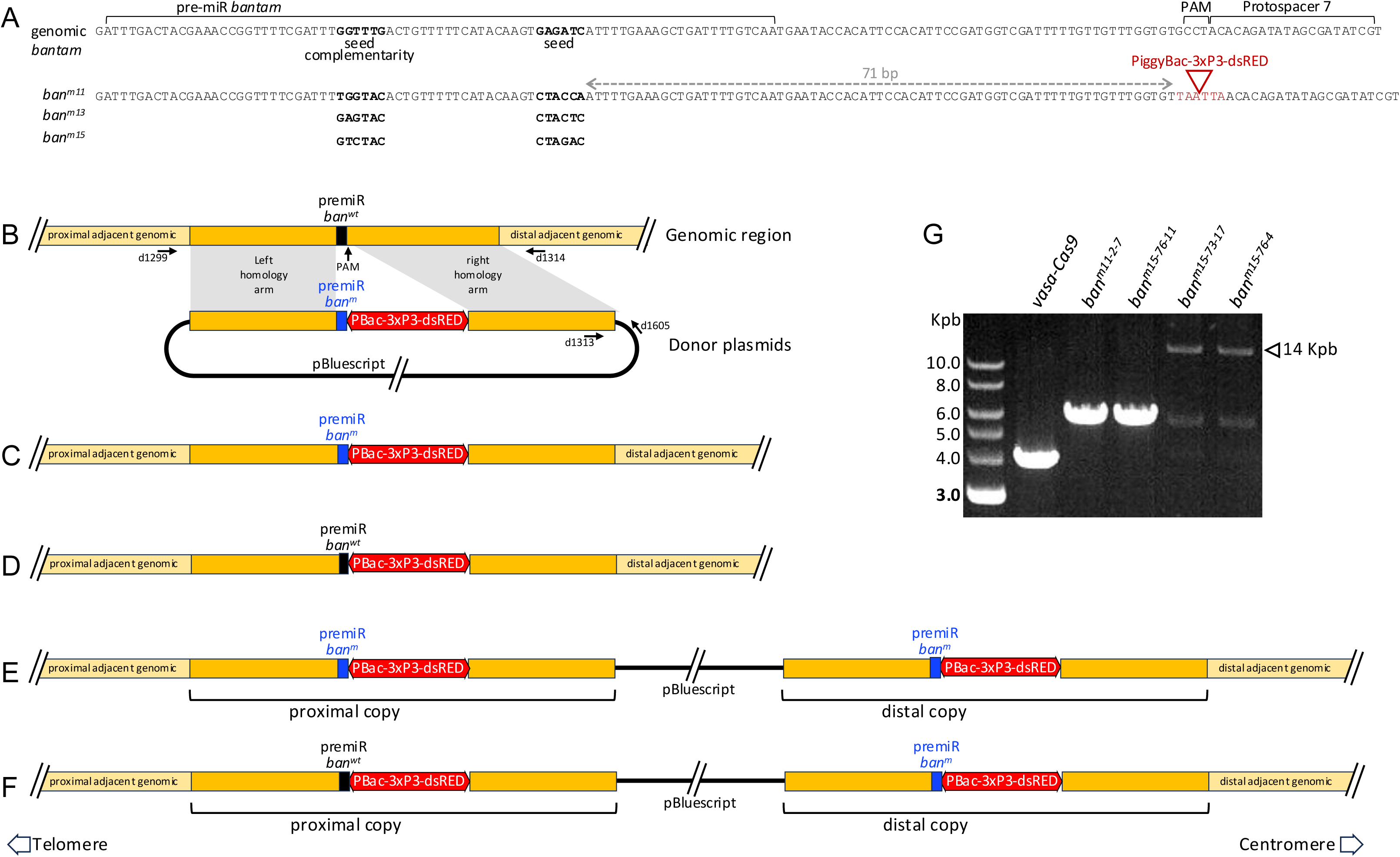
HDR-Scarless replacement of *bantam*. **A.** Sequences of the wild type *bantam* locus and three designed *bantam* variant alleles *ban^m11^*, *ban^m13^*, *ban^m15^*. Horizontal brackets above the *bantam* wild type sequence indicate the positions of the *bantam* pre-miR, protospacer and PAM used in the study. The positions of six nucleotides of the *bantam* wild type or mutated seed and of the upstream seed complementary sequences are indicated in bold. The site of integration of the *PiggyBac-3xP3-dsRED* cassette is indicated with a triangle, and the nucleotides that were inserted in the genomic DNA around the PAM are in red. The *PiggyBac-3xP3-dsRED* cassette contains *3×P3* eye-specific promoter and DsRed flanked by two inverted repeats TTAA to be recognized by the *PiggyBac* transposase. A grey dashed arrow indicates the 71 bp spacer between *bantam* seed and the cassette. **B-F.** Donor constructs design (**B**) and schematic organization of the *bantam* locus in edited lines, homozygous lethal (**C, E**) or viable (**D, F**), without (**C, D**) or with (**E, F**) plasmid integration. The upper part in (**B**) depicts the *bantam* genomic region, with a black box representing the wild type *bantam* pre-miR location and a vertical arrow showing the PAM position. The lower part schematizes the organization of the donor plasmids pBT-*ban^m11^-dsRED*, pBT-*ban^m13^-dsRED* and pBT-*ban^m15^-dsRED.* The thick black line represents the pBluescript vector. The blue box represent the mutated *bantam* pre-miR location. Thin arrows indicate the approximative positions of oligonucleotides d1299 and d1314 (for long-range PCR of the edited locus) and d1313 and d1605 (to amplify the junction between the right homology arm and the pBluescript vector). Homology arms present in the donor plasmids are indicated in orange, and the adjacent genomic DNA around the *bantam* locus but absent from the donor plasmids in in a lighter tone of orange. The *PiggyBac-3xP3-dsRED* cassette is shown as a red double arrow. Drawings are not to scale. **G.** Examples of PCR products generated with genomic DNA from some edited animals migrated on a 1% agarose gel stained with ethidium bromide. The empty arrowhead marks the position of a ∼14 Kbp PCR fragment in the two rightmost lanes. Lower bands in these two lanes (around 5.7 Kbp) likely correspond to contaminating PCR.

From these injections, 5, 13, and 12 fertile G0 adults from embryos injected with *ban^m11^*, *ban^m13^*, and *ban^m15^* donor plasmids, respectively, produced at least one fluorescent offspring (Table 1). In several cases, G0 adults yielded only a single fluorescent progeny (e.g., *ban^m11^*#110; *ban^m13^* #7, #19, #26, #36, #50; and *ban^m15^* #11, #82, #91, #107). Four of these fluorescent individuals (*ban^m13^* #26; *ban^m15^* #82, #91, #107) died before reproducing.

In many other cases, G0 adults produced multiple fluorescent offsprings, up to twenty-five in the case of *ban^m15^* #73 and twenty-eight in the case of *ban^m15^* #26 and #76. Initially, individual fluorescent animals were used to establish independent lines, which were subsequently assessed for homozygous viability or lethality using the *TM6B* balancer stock marked with *Sb* and *Tb*.

Surprisingly at first, siblings derived from a single G0 founder sometimes gave rise to both viable and lethal lines. The lethal lines displayed the characteristic *bantam* mutant phenotype. This phenomenon was observed for *ban^m11^* #15 and #104; *ban^m13^* #14, #69, #71, and #81; and *ban^m15^* #26, #31, #72, #73, and #76. These findings suggested that multiple distinct editing events might have occurred within the germline of individual G0 adults, which was confirmed thereafter.

### Various alterations produced by CRISPR-Cas9-HDR in the *bantam* locus

To characterize the molecular organization of the *bantam* locus in edited lines, homozygous animals were individually processed for DNA extraction followed by long-range PCR using primers mapping to genomic regions flanking the homology arms used in the donor plasmids (Figure 4B). In successfully edited alleles, this PCR was expected to produce a 5.7 kb fragment due to the insertion of the *PiggyBac-3xP3*-dsRED cassette. This was frequently observed (e.g., *ban^m11-2-7^* and *ban^m15-76-11^*, Figure 4G) in contrast to the 4 kb fragment product amplified from the wild-type *vasa*-*Cas9* strain. Subsequent Sanger sequencing of these PCR products revealed that those from lethal lines contained both the expected *ban^m11^*, *ban^m13^*, or *ban^m15^* mutation and the *PiggyBac-3xP3*-dsRED insertion. This confirmed that homologous recombination had fully replaced the endogenous *bantam* allele with the modified donor sequence, as intended (Figure 4C). In contrast, PCR products of the same expected 5.7 kb size obtained from viable lines did not contain the engineered mutations, but, instead, had retained the wild-type *bantam* sequence (Figure 4D). This suggested that, in these cases, recombination may have occurred downstream of the targeted mutation, possibly within the 71 bp spacer between the mutation site and the PiggyBac cassette (Figure 4A), rather than within the designated left homology arm. As a result, only part of the donor sequence was incorporated, leaving unaltered the sequence of the mature *bantam* miRNA despite successful marker integration.

In some lines, the long-range PCR failed to efficiently amplify the expected 5.7 kb fragment. Instead, a faint band of roughly 14 kb was observed (e.g., *ban^m15-73-17^* and *ban^m15-76-4^*, Figure 4G), suggesting the presence of a complex genomic rearrangement. These situations were detected in a total of nine lines: five *ban^m13^* (two lethal and three viable) and four *ban^m15^* (two lethal and two viable). To investigate whether the entire donor plasmid had been inserted at the target site in these lines, we performed PCR to amplify the junction between the plasmid backbone and the right homology arm (Figure 4B). Successful amplification of this junction confirmed that the full donor plasmid was indeed integrated into the genome in the nine lines, possibly at the Cas9-induced cleavage site. Such integrations should result in tandem duplication of the *bantam* locus, as illustrated in Figure 4E and F. Subsequently, to characterize the organization of the *bantam* locus in these lines, the proximal and distal duplications of the *bantam* region were amplified and sequenced separately. In all cases, the distal copy (respective to the chromosome telomere) of *bantam,* originating from the donor plasmid, contained the engineered mutation. The proximal copy carried either the mutated *bantam* allele (in lethal lines, Figure 4E) or the wild-type allele (in viable lines, Figure 4F). Interestingly, both lethal and viable lines consistently harboured the *PiggyBac-3xP3*-dsRED cassette within the proximal copy. This indicated that homology-directed repair (HDR) occurred at the proximal region in all cases, in parallel with full-length integration of the donor plasmid. Thus, the resulting configuration involved both HDR and plasmid insertion, producing a duplicated *bantam* locus with distinct allele compositions.

In conclusion, the CRISPR-Cas9-HDR experiments described above successfully established stable lines containing the dsRED marker, representing four distinct outcomes: (1) introduction of the *bantam* mutation (Figure 4C), (2) no modification of the wild-type *bantam* allele (Figure 4D), (3) donor plasmid integration resulting in locus duplication with the *bantam* mutation (Figure 4E), and (4) donor plasmid integration resulting in locus duplication without the *bantam* mutation (Figure 4F). These outcomes frequently occurred independently within the germline of a single G0 adult. Notably, one embryo injected with the *ban^m15^* donor plasmid gave rise to independent events corresponding to all four situations (Table 1, G0 #76; lines *ban^m15-76-4^, ban^m15-76-11^, ban^m15-76-16^*, and *ban^m15-76-23^*).

### Does the *PiggyBac-3xP3*-dsRED cassette interact with *bantam* function ?

The scarless allele replacement method proved highly efficient for generating precise *bantam* gene variants via CRISPR-Cas9-HDR. As described above, the experiments successfully produced three, eight, and eight independent lines carrying the expected *ban^m11^*, *ban^m13^*, and *ban^m15^* mutations, respectively, along with the dsRED marker (Table 2). For each mutation, two stable lines were selected for excision of the *PiggyBac-3xP3*-dsRED cassette using the PiggyBac transposase (Table 1). The resulting reconstituted lines, which retained the *ban^m11^*, *ban^m13^*, or *ban^m15^*mutations but lacked the dsRED cassette, continued to exhibit the characteristic *bantam* mutant phenotypes. These findings indicated that the *PiggyBac-3xP3*-dsRED cassette, positioned 48 nucleotides downstream of the *bantam* pri-miR, has minimal, if any, impact on *bantam* function.

**Table 2:** Categories of edits generated by CRISPR-Cas9-HDR.

|  | Number of independent edits (%) |  |  |
| --- | --- | --- | --- |
|  | *banm11* | *banm13* | *banm15* |
| **Mutated *bantam*** | 3 (43%) | 8 (40%) | 8 (50%) |
| **Unmutated *bantam*** | 4 (57%) | 7 (37%) | 4 (25%) |
| **Donor integration** | 0 (0%) | 5 (25%) | 4 (25%) |
| **Total** | 7 | 20 | 16 |

In addition, a substantial number of lines were recovered in which the sequence of the mature *bantam* miRNA remained unaltered despite successful integration of the fluorescent dsRED marker—four, seven, and four such lines for *ban^m11^*, *ban^m13^*, and *ban^m15^*, respectively. We did not detect phenotypical differences between these viable, unmutated lines and wild-type flies, aside from their fluorescent eyes. This further confirms that the *PiggyBac-3xP3*-dsRED insertion at this downstream position is phenotypically neutral. These unmodified *bantam* lines carrying the dsRED cassette serve as ideal controls for the *ban^m11^*, *ban^m13^*, and *ban^m15^* mutant lines that still retain the cassette.

The overall conclusion is that, despite many unwanted outcomes, the scarless allele replacement method was very efficient in our hands to produce precise *bantam* gene variants using CRISPR-Cas9-HDR.

## DISCUSSION

The outcomes of CRISPR-Cas9 editing experiments are influenced not only by the targeted site and experimental design, but also by the organism and tissue in which they are performed (Meyenberg et al., 2021). Therefore, gaining a comprehensive and in-depth understanding of the editing landscape in specific contexts provides invaluable insights for future gene-editing applications in similar conditions. In this paper, we present the results of CRISPR-Cas9 experiments utilizing either the NHEJ or HDR repair pathways to generate germline mutations affecting the *bantam* miRNA in *Drosophila*. By analysing a large collection of edited lines at the molecular level, we reveal a diverse and somewhat unexpected range of outcomes produced by each pathway under these conditions.

Both approaches were notably efficient, presumably primarily due to the initial Cas9-mediated cleavage events, which depend heavily on guide RNA design. In these studies, options were limited: few protospacers were predicted near the *bantam* locus. The guide RNAs that we selected did not fully meet the optimal design criteria (Ren et al., 2014; Haussmann et al. 2024). However they had no predicted off-targets in the genome version available at the time, and they proved to be sufficiently effective for our experimental goals.

### NHEJ Predominantly Acts Independently at Two Close Cleavage Sites

The experiments utilizing the NHEJ pathway were designed with two guide RNAs targeting the *bantam* locus 90 bp apart, aiming to create a deletion that would remove most of it. This would provide a rationale for adapting this strategy to efficiently inactivate *bantam* in tissues beyond the germline (Meltzer et al., 2019; Port et al., 2020; Port and Boutros, 2022).

Contrary to our expectations, deletions spanning both target sites were rare, and only one exhibited breakpoints precisely at the predicted Cas9 cleavage sites. Nonetheless, most edited lines contained indels at one or both target sites, with the majority of these alterations disrupting *bantam* function. The editing patterns we observed were consistent with previous studies (Gratz et al., 2014; Port et al., 2014; Yu et al., 2013), typically involving small, independent events at each site. These findings suggest that editing occurs independently at closely spaced target sites, an insight that may inform the design of future experiments aimed at introducing simultaneous mutations at multiple loci.

Notably, we did not recover any lines in which the *bantam* locus remained completely unedited. Even in viable lines where *bantam* function was preserved, sequence alterations were present: specifically, a single cytosine deletion at the proximal target site. The absence of unmodified target sequences does not necessarily indicate that precise repair never occurred. Rather, imprecise repair events may have disrupted the sgRNA recognition site, preventing further Cas9 activity. In contrast, precisely repaired sites would remain susceptible to repeated cleavage, increasing the likelihood of subsequent imprecise modifications. Consequently, imprecise repair events would be expected to accumulate over time, ultimately outnumbering precise ones.

A particularly noteworthy outcome was the recovery of a hypomorphic *bantam* allele, *ban^d1-44^*, which supports viability but exhibits clear growth defects. This allele presents a valuable opportunity to dissect the distinct roles of *bantam* in viability and growth regulation. In the short term, we will focus on characterizing its expression dynamics, maturation profile, and RNA products. Over the longer term, *ban^d1-44^* will serve as a powerful tool for investigating functional interactions between *bantam* and its downstream targets.

### Effectiveness of the HDR Pathway

HDR-based experiments were aimed to replace the wild-type *bantam* locus with the mutant variants *ban^m11^, ban^m13^, ban^m15^*, each linked to a fluorescent marker to facilitate screening. These experiments were successful as multiple independent lines carrying each mutation were established. This outcome underscores the remarkable efficiency of the HDR pathway using an appropriate donor over NHEJ for generating precise, targeted deletions using CRISPR-Cas9.

Despite the overall efficiency, we also observed outcomes that were unexpected to us, falling into two main categories: partial replacements, where the *PiggyBac-3xP3*-dsRED cassette was integrated without the intended *bantam* modification, and full integrations of the donor plasmid into the genome. In several cases multiple independent editing events occurred simultaneously in the germ cells of one founder.

### Unexpected HDR Outcomes: Partial Homology-Directed Replacements

Unmutated versions of *bantam* were recovered at high frequencies, comparable to the mutated versions: 57 vs. 43%, 37 vs. 40%, and 25 vs. 50% for *ban^m11^, ban^m13^,* and *ban^m15^*, respectively (Table 2). These mutants carried the *PiggyBac-3xP3*-dsRED cassette but lacked the engineered portion of the *bantam* locus, indicating that the proximal breakpoint for homology-directed repair (HDR) was located downstream of *bantam*.

This outcome was unexpected, as we initially assumed that the likelihood of recombination occurring within the small 71 bp spacer sequence between *bantam* and the predicted cleavage site would be lower than anywhere else in the 1.2 kb left homology arm. Indeed, a previous study established a 1 kb threshold for efficient HDR using donor plasmids following induced DNA breaks in *Drosophila* (Beumer et al., 2013). In contrast, much shorter homology arms—even less than 100 nucleotides—were reported to be sufficient when using donors in the form of linear DNA (Gratz et al., 2013; Kanca et al., 2019, 2021; Bui and Kamiyama, 2023). Our findings suggest that short homology arms in donor plasmids may still support efficient HDR. Nonetheless, it is plausible that some degree of homologous pairing between the donor plasmid and the genomic locus to be edited could be a prerequisite for HDR. This pairing would be favoured by longer homology arms, even if the homology is locally disrupted by the seed and complementary sequences, as observed in our study.

Thus, despite the careful design of our CRISPR-Cas9 HDR experiment, we failed to anticipate a frequent outcome: partial repair due to a short homology sequence between the fluorescent marker and the intended mutation. As was alluded to earlier, these unmutated versions of *bantam* serve as ideal controls for the *ban^m11^, ban^m13^,* and *ban^m15^* mutant lines. Consequently, we consider them to be highly valuable and positive outcomes of our strategy.

### Unexpected HDR Outcomes: Donor Plasmid Integrations

In experiments utilizing the HDR pathway to generate the *ban^m13^* and *ban^m15^* mutations, we occasionally observed the complete integration of the donor plasmid. These integrations resulted in the duplication of the *bantam* locus, with the presumed initial genomic copy being separated from the plasmid-derived copy by vector sequences. These integrations occurred alongside other editing events: in all 10 studied cases, the presumed initial *bantam* copy was edited to contain the *PiggyBac-3xP3*-dsRED cassette, and in four cases, the intended *ban^m13^* or *ban^m15^* mutations were also successfully introduced.

Unlike the unmutated *bantam* alleles described earlier, these plasmid integrations are not useful for our experiments. More concerningly, they were initially overlooked, as they were not detected by standard PCR analyses designed to verify the expected editing outcomes. Instead, they were identified through long-range PCR, which captured the full genomic region beyond the donor plasmid fragment.

Unintended template DNA integrations are a known phenomenon in genome editing, including CRISPR-Cas9 and other techniques such as TALEN. These events are thought to result from homology-independent targeted integration (Suzuki et al., 2016). They have been documented in various organisms, including nematode (Dickinson et al., 2013), zebrafish (Stemmer et al., 2015), calves (Carlson et al., 2016; Norris et al., 2020) , mice (Skryabin et al., 2020; Zhao et al., 2024), as well as in many experiments in cultured cells including mammalian embryonic stem cells (Bi et al., 2025; Erbs et al., 2023; Hendel et al., 2014; Liu et al., 2021). This issue is particularly critical in therapeutic applications, where unintended genomic alterations are unacceptable (Zhao et al., 2024). As a result, several researchers have proposed strategies to maximize homology-dependent repair (HDR) and minimize unwanted integrations (Hermantara et al., 2024; Karasu et al., 2024; Zhang et al., 2017). Recently, in *Drosophila*, an elegant system based on the single-strand annealing (SSA) pathway was described (Aguilar et al., 2024). However, its implementation relies on two successful recombination events, which makes the process relatively complex and time-consuming.

In basic research involving model organisms like *Drosophila*, unintended template DNA integrations may not always be considered a major issue. However, it is essential to be aware of these occurrences, systematically identify them, and discard them as early as possible to prevent misinterpretation of results. The work presented here provide some strategic plans to this goal.

## Data Availability Statement

Fly lines and plasmids are available upon request. Annotated sequences *will be available at* https://github.com *– in progress*.

File S1 contains supplemental Tables S1-5.

File S2 contains sequences related to Figure 2 in fasta format.

File S3 contains supplemental Figures S1 and S2.

## Acknowledgments

We acknowledge the imaging facility MRI, member of the national infrastructure France-BioImaging (https://ror.org/01y7vt929) supported by the French National Research Agency (ANR-24-INBS-0005 FBI BIOGEN), as well as the Biocampus Drosophila facility. Stocks obtained from the Bloomington Drosophila Stock Center (NIH P40OD018537) were used in this study. Our work is supported by the CNRS, University of Montpellier, and Labex EpiGenMed.

